# OsUVR8b, rather than OsUVR8a, plays a predominant role in rice UVR8-mediated UV-B response

**DOI:** 10.1101/2023.12.07.570679

**Authors:** Yu-long Chen, You-bin Zhong, David W. M. Leung, Xiao-Yu Yan, Meng-ni Ouyang, Yu-zhen Ye, Shi-mei Li, Xin-xiang Peng, E-e Liu

**Author notes:** **Mailing address and email address of one author for contact: Name**: E-e Liu, **Address**: College of Life Sciences, South China Agricultural University, Guangzhou, 510642, P.R. China, **E-mail address** or. contributed equally to Yu-long Chen.

## Abstract

UV RESISTANCE LOCUS 8 (UVR8) has been identified in *Arabidopsis thaliana* as the receptor for UV-B radiation mediating photomorphogenic responses and acclimation to UV-B radiation. However, UVR8-mediated UV-B signaling pathways in rice, that has two proteins (UVR8a and UVR8b) with homology to AtUVR8, remain largely unknown. In this study, *UVR8a* and *UVR8b* were found to be expressed mainly in rice leaves and leaf sheaths, while the level of UVR8b was higher than that of UVR8a. In agreement with prior studies on AtUVR8, *uvr8b* and *uvr8a uvr8b* rice mutants exposed to UV-B showed reduced UVB-induced growth inhibition and upregulation of *CHS* and *HY5* transcripts along with acclimation to UV-B, overexpressing UVR8a or UVR8b enhanced UV-B-induced growth inhibition and acclimation to UV-B, compared to wild-type plants. UV-B was able to enhance the interaction between CONSTITUTIVE PHOTOMORPHOGENESIS1 (COP1) with UVR8a/UVR8b, whereas the interaction intensity of REPRESSOR OF UV-B PHOTOMORPHOGENESIS2 (RUP2) with UVR8a was significantly higher than that with UVR8b. In addition, UVR8a and UVR8b were also found in the nucleus and cytoplasm, but OsUVR8 proteins were localized in nucleus in the absence of UV-B. The level of OsUVR8 monomer showed an invisible change in the leaves of rice seedlings transferred from white light to white light supplemented with UV-B, even UV-B can weaken the interactions of UVR8a or/and UVR8b. Therefore, both UVR8a and UVR8b, that have different location and response modes with Arabidopsis UVR8, function in the response of rice to UV-B radiation, whereas UVR8b plays a predominant role in this process.

## Introduction

Approximately 95% of UV-B radiation is absorbed by the ozone layer and the rest reaches the Earth’s surface with an average intensity of 100 μW. cm^−2^(Cejka et al, 2011). Therefore, exposure to UV-B radiation is inevitable for higher plants when grown under sunlight because they are sessile. Reduced biomass and inhibition of hypocotyl elongation are the most common morphological effects following exposure to UV-B radiation (Vandenbussche et al., 2018). In addition, reduced pollen fertility in various plant species has been observed in the response to UV-B radiation (Jansen et al., 1998; Caldwell et al., 2003). In general, plants respond differently to high or low doses of UV-B irradiation. High-level UV-B radiation induces DNA damage, generates reactive oxygen species (ROS) and impairs photosynthesis (Chen et al, 2022; Frohnmeyer and Staiger, 2003). In contrast, non-damaging low doses of UV-B radiation (<1 μmol m^−2^ s^−1^) can act as a regulatory signal that is specifically perceived by the UV-B photoreceptor, UV RESISTANCE LOCUS 8 (UVR8). In response to UV-B exposure, plant photomorphogenesis and acclimation to UV-B is triggered by stimulating protective mechanisms or activating repair mechanisms to cope with UV-B radiation (Brown et al, 2005). The protective action against UV-B radiation is based on UVR8-mediated expression of various genes, including those encoding DNA photorepair enzymes and flavonoid synthesis enzymes that provide a UV-B-absorbing sunscreen in the epidermal tissues. Upon UV-B irradiation, the UVR8 photoreceptor is converted from a biologically inactive dimer to two active monomers which interact with COP1, a ubiquitin ligase, to prevent poly-ubiquitination of the transcriptional factor HY5. Consequently, as the degradation of HY5 is suppressed and the photomorphogenic response can then be initiated (Favory et al., 2009; Cloix et al., 2012). Acting downstream of the specific UVR8-mediated UV-B protective responses, HY5 induces the synthesis of the REPRESSOR UV-B PHOTOMORPHOGENESIS PROTEIN 1 (RUP1) and RUP2, which interact with the UVR8 monomers to release COP1. This would result in the re-dimerization of UVR8 monomers and repression of the UVR8-mediated UV-B signaling (Gruber et al., 2010; Heijde and Ulm, 2013). This induced UV-B acclimation response and negative feedback balancing mechanism have been thoroughly characterized in Arabidopsis, where studies have already shown that the AtUVR8 receptor is a key to UV-B acclimation response. Relatively little is, however, known about the underlying mechanisms of UV-B perception and signal transduction in other plant species.

Although Arabidopsis is an important model experimental plant, many UV-B acclimation mechanisms described in this species maybe different from those in other plants. For instance, in some plants, there are proteins with low homology to AtRUP1 and AtRUP2, and two proteins with homology to AtUVR8 (Tossi et al, 2019). In rice, there is only one protein with 46% homology to AtRUP2 and two proteins OsUVR8a (Os02g0554100) and OsUVR8b (Os04g0435700) that are homologous to AtUVR8 (Figure S1A). Since rice is adapted to growth in a high light environment with high levels of UV-B, the mechanism of UV-B acclimation in rice may therefore be different from that of Arabidopsis. However, remarkably little is known about the underlying mechanisms of the UV-B perception and signal transduction in rice. Despite UV-B signaling in rice has been reported (Idris et al, 2021), whether OsUVR8a and OsUVR8b function as the UV-B photoreceptor in rice is still unclear. In this work, the UV-B response mediated by UVR8a and UVR8b in rice was investigated, particularly in comparison with that in Arabidopsis. The findings here suggest that OsUVR8b, rather than OsUVR8a, plays a predominant role in UV-B acclimation. Besides, the mechanisms of UVR8-mediated UV-B perception and signal transduction in rice might be different from that in Arabidopsis although UVR8 is thought to be conserved among different plants.

## Results

### The abundance of *OsUVR8b* is higher than that of *OsUVR8a* in rice seedlings

The differences in the amino acid sequences and tertiary structures among OsUVR8a, OsUVR8b and AtUVR8 major in N- and C-termini. The tertiary structure of OsUVR8a is more similar to that of AtUVR8 than that of OsUVR8b (Figure S1). Since the C-terminus plays an important role in UV-B signaling, the differences in amino acid sequences at the C-termini suggest that there might be some differences in their participation in UV-B signaling. Moreover, there are significant differences in their promoter sequence, suggesting that their regulation may be different. The transcript levels of *OsUVR8a* and *OsUVR8b* were much higher in rice leaves and leaf sheaths than in roots. In these three plant parts, the levels of *OsUVR8b* transcripts were higher than those of *OsUVR8a* (Figure 1A). Correspondingly, the levels of UVR8b protein in these tissues was also higher than those of UVR8a protein, but that of UVR8 protein in leaf sheaths was much higher than that in leaves and roots (Figure 1C). The transcript levels of *UVR8a* and *UVR8b* were downregulated in the leaves of rice seedlings transferred from white light to white light supplemented with UV-B (Figure 1B). However, the OsUVR8 protein level did not seem to change (Figure 1D) in the leaves of rice seedlings exposed to UV-B while the level of the OsUVR8 dimer slightly decreased accompanied by an increased level of OsUVR8 monomer (Figure 1D). Whereas the monomers of AtUVR8 increased significantly in the leaves of Arabidopsis exposed to same UV-B lamp (Figure S2). Moreover, the levels of *HY5*, *RUP2* and *CHS* transcripts increased significantly in the leaves of rice seedlings transferred from white light minus UV-B to that supplemented with UV-B (Figure S3).

**Figure 1.**
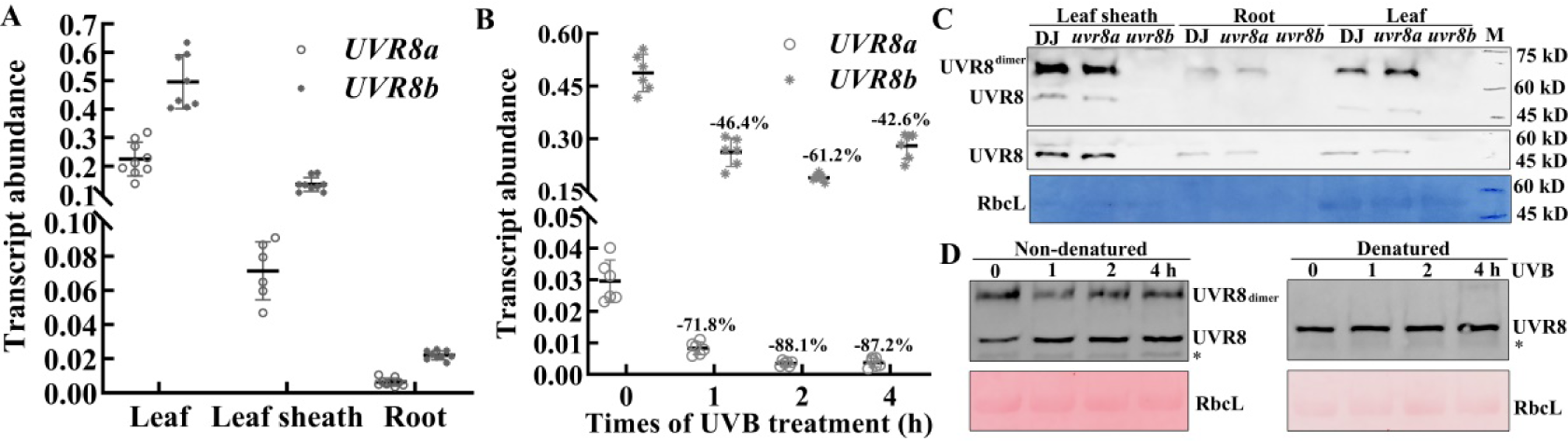
Abundance of OsUVR8b is higher than that of OsUVR8a in rice seedlings. A: qRT-PCR analyses of *OsUVR8a* and *OsUVR8b* transcripts in various tissues of 5-leaf stage wild-type (WT) rice seedlings (cv. Donjin or DJ) grown in the glasshouse. B: qRT-PCR analyses of *OsUVR8a* and *OsUVR8b* transcripts in the leaves of 2-leaf stage WT rice seedlings grown in a growth chamber under 200-750 µmol·m^−2^·s^−1^ white light with or without supplementation of 75-85 μW·cm^−2^ UV-B radiation for 0, 1, 2 and 4 h in a growth chamber, respectively. C: immunoblot analysis of UVR8 protein levels in the leaves (20 μg protein/lane), leaf sheaths (10 μg protein/lane) and roots (20 μg protein/lane) of 4-leaf stage WT, *uvr8a* and *uvr8b* seedlings grown under sunlight, amido black staining is shown as a loading control. D: immunoblot analysis of UVR8 protein in the leaves of 3-leaf stage WT rice seedlings grown in a growth chamber under 200-750 µmol·m^−2^·s^−1^ white light or without supplementation of 70-80 μW·cm^−2^ UV-B radiation for 0, 1, 2 and 4 h. Ponceau staining of ribulose-1, 5-bisphosphate carboxylase/oxygenase (Rubisco) large subunit (RbcL) is shown as a loading control.

### UVR8b plays a predominant role in UVR8-mediated UV-B responses

To investigate any differences between UVR8a and UVR8b in mediating UV-B response, the following *uvr8a*, *uvr8b* and *uvr8a uvr8b* mutants were generated using the CRISPR/Cas9 techniques (Figure S4). The UVR8 protein levels in the leaves of *uvr8b* was remarkably lower than that in *uvr8a*, while no UVR8 protein was detectable in the leaves of *uvr8a uvr8b* (Figure 2A). Compared to seedlings of WT, *uvr8a* and *uvr8b* mutants grown under white light, those grown under specific UV-B irradiation (50-100 μW·cm^−2^ UV-B) for 7 days exhibited significantly lower (*p*<0.05) plant height (Figure 2B and C). However, the growth inhibition in *uvr8b* grown under white light with UV-B compared to that grown under light without UV-B mutants was relatively smaller than that in WT and *uvr8a* under the same experimental conditions (Figure 2B and C). There was no significant difference in the growth of *uvr8a uvr8b* grown under white light with or without narrowband UV-B supplementation (Figure 2B and 2C). Two-leaf stage rice seedlings were grown under white light without UV-B supplementation (non-acclimation treatment) or exposed to 50-100 μW.cm^−2^ UV-B (acclimation pretreatment) for 8 days in the incubator before both groups of seedlings were exposed to 90-110 μW.cm^−2^ UV-B for 0 and 4 h and were then returned to recovery under white light without UV-B supplementation for 48 h (Figure 2D). In the non-acclimation treatment, obvious lesions appeared in the leaves of WT and the mutants exposed to the high UV-B for 4 h (Figure 2D), but relatively more prominent lesions were only displayed in the leaves of *uvr8a uvr8b* mutants grown under white light supplemented with specific UV-B in the incubator and then exposed to UV-B for 4 h. Visible lesions were not found in the leaves of WT in the acclimation treatment (Figure 2D). The plant height of *uvr8a uvr8b* grown under sunlight in the days with a medium or lower UV-B level was also higher than that in WT and *uvr8a* mutants, suggesting UVR8a and UVR8b are both involved in UV-B-induced growth inhibition(Figure S5A). During sunny day in the hot summer in Guangzhou (from June to September), China, there were no significant differences in the plant height of *uvr8a uvr8b*, *uvr8a* and WT seedlings (Figure S5B and C). However, significant lesion mimic appeared in leaves of *uvr8a uvr8b* seedlings grown under sunlight from June to August (Figure S5D), and only fewer lesion mimic found in leaves of *uvr8b* seedlings and much less in the leaves of *uvr8a* seedling (Figure S5D), suggesting that the tolerance of *uvr8b* and *uvr8a uvr8b* seedlingsto UV-B was more reduced. Consistent with the reduced UV-B tolerance phenotype of *uvr8a uvr8b* and *uvr8b*, the transcript levels of the well-characterized genes of chalcone synthase (*CHS*) and *HY5* activated by UV-B were lower in *uvr8b* than in WT, while *uvr8a* mutants had similar *CHS* and *HY5* transcript levels compared with WT after exposure to UV-B, *uvr8a uvr8b* mutants exhibited significant reduced levels of *CHS* and *HY5* (Figure 2E and 2F). Moreover, *uvr8a* mutants exhibited slightly reduced seed production rate and plant height as well as comparable pollen vitality compared with WT grown in a paddy field, while plant height, seed production rate and pollen vitality of *uvr8b* mutants were significantly lower than that of WT (Fig S5E and H), *uvr8a uvr8b* mutants also exhibited significant decrease in seed production rate and pollen vitality compared with WT, further supporting that UVR8b plays a major role in UV-B-induced photomorphogenesis and rice tolerance to UV-B.

**Figure 2.**
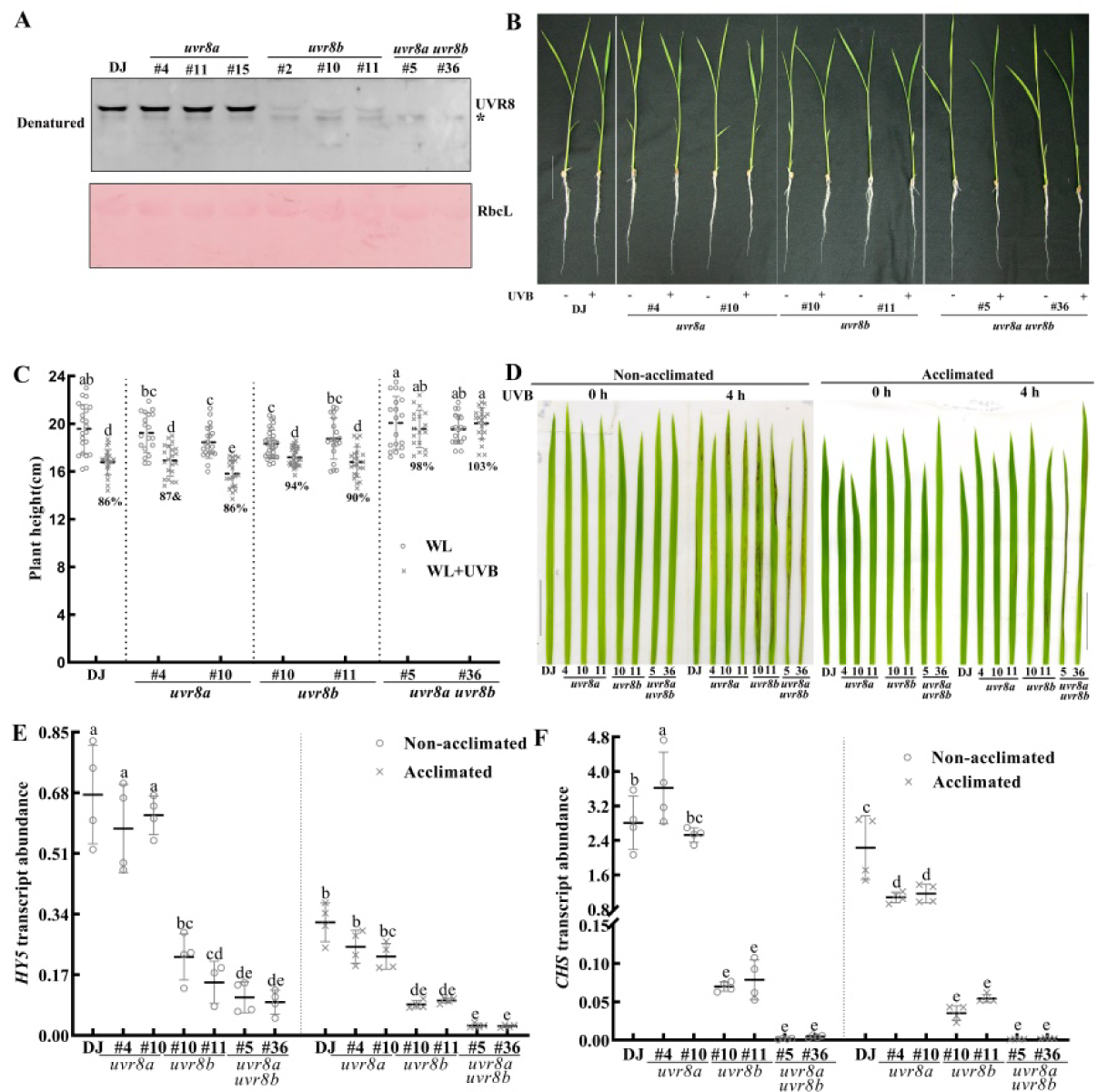
Mutation of *UVR8a* or/and *UVR8b* impairs the response and acclimation to UV-B of rice seedlings. A: immunoblot analysis of UVR8 protein levels in the leaves (second leaves from top to bottom) of 5-leaf stage wild type (DJ), *uvr8a* (#4, #11, #15), *uvr8b* (#2, #10, #11) and *uvr8a uvr8b* (#5, #36) seedlings grown in a paddy field, Ponceau staining of RbcL is shown as a loading control. B: phenotype of representative seedlings grown under white light for 2 d after germination and then to white light or white light supplemented with 50-100 μW·cm^−2^ UV-B for 7 d. C: quantification of plant height of the seedlings depicted in B. D: phenotype of seedlings grown under white light for 2 d after germination and then grown for 8 d in white light (non-acclimated) or white light supplemented with 50-100 μW·cm^−2^ narrowband UV-B (acclimated) and then exposed to 90-110 μW·cm^−2^ UV-B stress for 0 and 4 h. Pictures were taken after a 2-d recovery period. E and F are quantitative RT-PCR analysis of *HY5* (E) and *CHS* (F) gene activation in response to UV-B in *uvr8a* (#4, #10), *uvr8b* (#10, #11) and *uvr8a uvr8b* (*#*5, *#*36) seedlings grown for 7 d in white light (non-acclimated) or white light supplemented with 50-100 μW·cm^−2^ UV-B (acclimated) and exposed to white light supplemented with 80-100 μW·cm^−2^ UV-B for 4 h compared to WT (DJ). The means assigned with the same letters indicate no statistically significant difference as determined by one-way different ANOVA with Tukey HSD or Duncan post-hoc multiple comparisons for differences among the means (*P* > 0.05).

### OsUVR8a and OsUVR8b were localized in nucleus even in the absence of UV-B

To determine the subcellular localization of OsUVR8a and OsUVR8b, the pYL322-*OsUVR8a*-GFP and pYL322-*OsUVR8b*-GFP fusion constructs were transformed into protoplasts from rice seedlings grown under sunlight. Expression of the OsUVR8–GFP fusion protein, like that of AtUVR8–GFP fusion protein, resulted in the GFP signal being detected in both the cytosol and nucleus, the GFP signal in the nucleus was well overlapped with that expression of the nucleus marker 35S:: OsABF1–RFP. In contrast, 35S::GFP showed a diffuse distribution in both cytosol and nucleus (Figure 3), suggesting that OsUVR8a and OsUVR8b is a nuclear- and cytosol-localized protein. Bright GFP fluorescence can still be observed in the nuclei even though expression of the OsUVR8–GFP fusion protein in protoplasts isolated from etiolated rice seedlings or *N. benthamiana* (transformed with GFP–OsUVR8) grown under white light (Figure S6), indicating OsUVR8 proteins still exist in nucleus in absence of UV-B.

**Figure 3.**
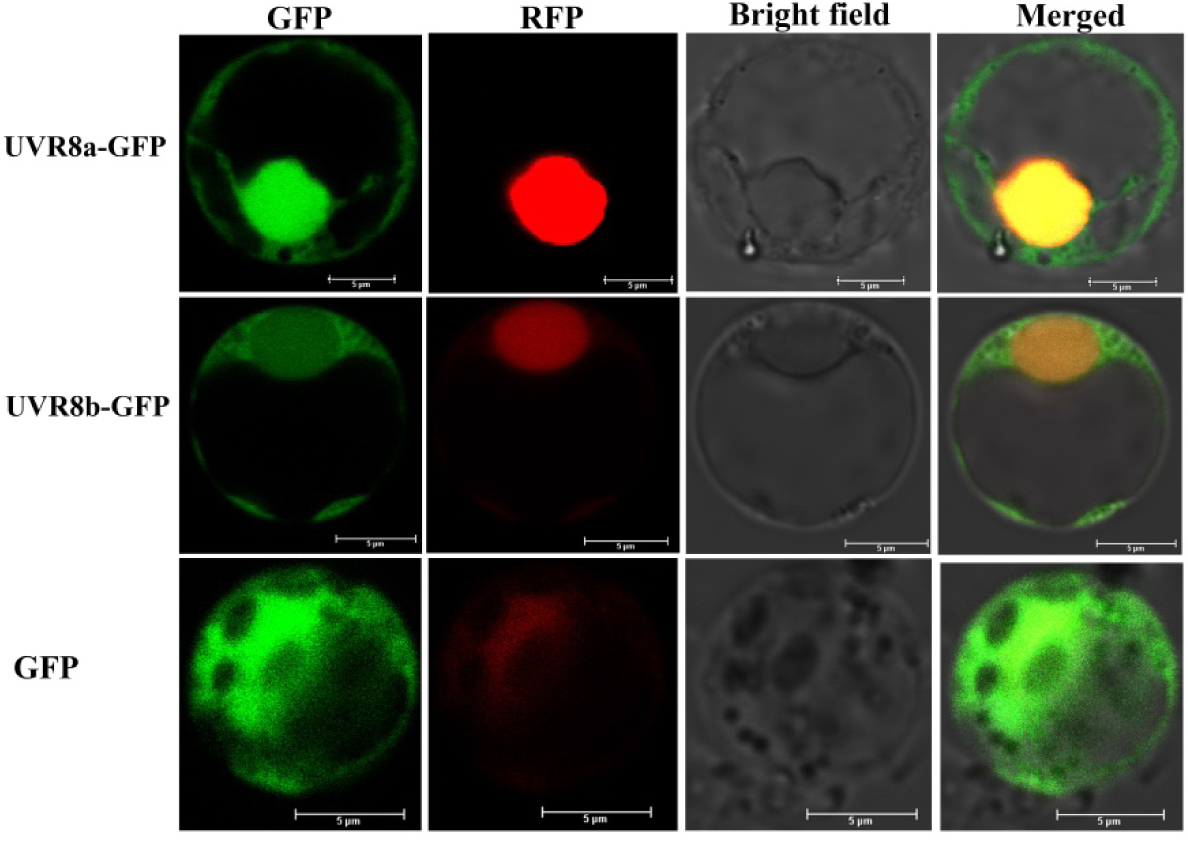
Fluorescent protein–tagged UVR8a and UVR8b proteins show nuclear and cytosolic localizations. Confocal micrographs of protoplasts were isolated from rice seedlings exposed to sunlight. The isolated protoplasts were co-transformed with the vectors pYL322-OsUVR8a-GFP (upper) and pYL322-OsABF1-RFP, and pYL322-OsUVR8b-GFP and pYL322-OsABF1-RFP (lower), respectively. OsABF1-RFP is a nuclear marker and GFP alone was used as a control. Bars=5 μm.

### OsUVR8a interacts with OsUVR8b, but UVR8a shows higher binding affinity to RUP2 than UVR8b

Tobacco leaves co-transformed with UVR8a-nLUC/UVR8b-nLUC and cLUC-UVR8a, cLUC-UVR8b, cLUC-COP1 and cLUC-RUP2 were grown under white light with or without supplementation with 230-250 μW·cm^−2^ UVB for 30 min after growing in the normal environment (under white light) for 36 h. The firefly luciferase substrate was then injected into the tobacco leaves before examining LUC activity using the NightSHADE LB98 Imaging System. The results showed that UVR8a interacted with UVR8b and UV-B was able to impair but not abolish completely their interaction, even when the tobacco leaves were exposed to 230-250 μW·cm^−2^ UV-B for 30 min (Figure 4A). In addition, Co-IP assay of rice endogenous UVR8 with UVR8-FLAG from 2-leaf P_UVR8b_: *UVR8-FLAG*/*uvr8b* seedlings transferred from white light (0 μW·cm^2^ UV-B) to sunlight also showed UVR8a interacted with UVR8b in sunlight (Figure 4D). Interestingly, fluorescence signals of OsUVR8a-nLUC and cLUC-OsUVR8a along with OsUVR8b-nLUC and cLUC-OsUVR8b were strong in firefly luciferase complementation experiments even when the tobacco plants were exposed to UV-B. However, tobacco transfected with AtUVR8-nLUC and cLUC-AtUVR8 showed no significant fluorescence signal, suggesting OsUVR8a and OsUVR8b might form dimer more easily than AtUVR8. When tobacco was placed under white light with 230-250 μW·cm^−2^ UV-B for 30 min, the interactions between COP1 and UVR8a / UVR8b were enhanced to different degrees, suggesting that UV-B could affect the interaction between COP1 and UVR8a / UVR8b (Figure 4B). The interaction between RUP2 with UVR8a / UVR8b occurred after white light and UV-B treatment, and the interaction of RUP2 with UVR8a was significantly stronger than UVR8b, suggesting that the interaction of RUP2 with UVR8a and UVR8b is different, and that the roles of UVR8a and UVR8b may be different in the response of rice to UV-B (Figure 4C).

**Figure 4.**
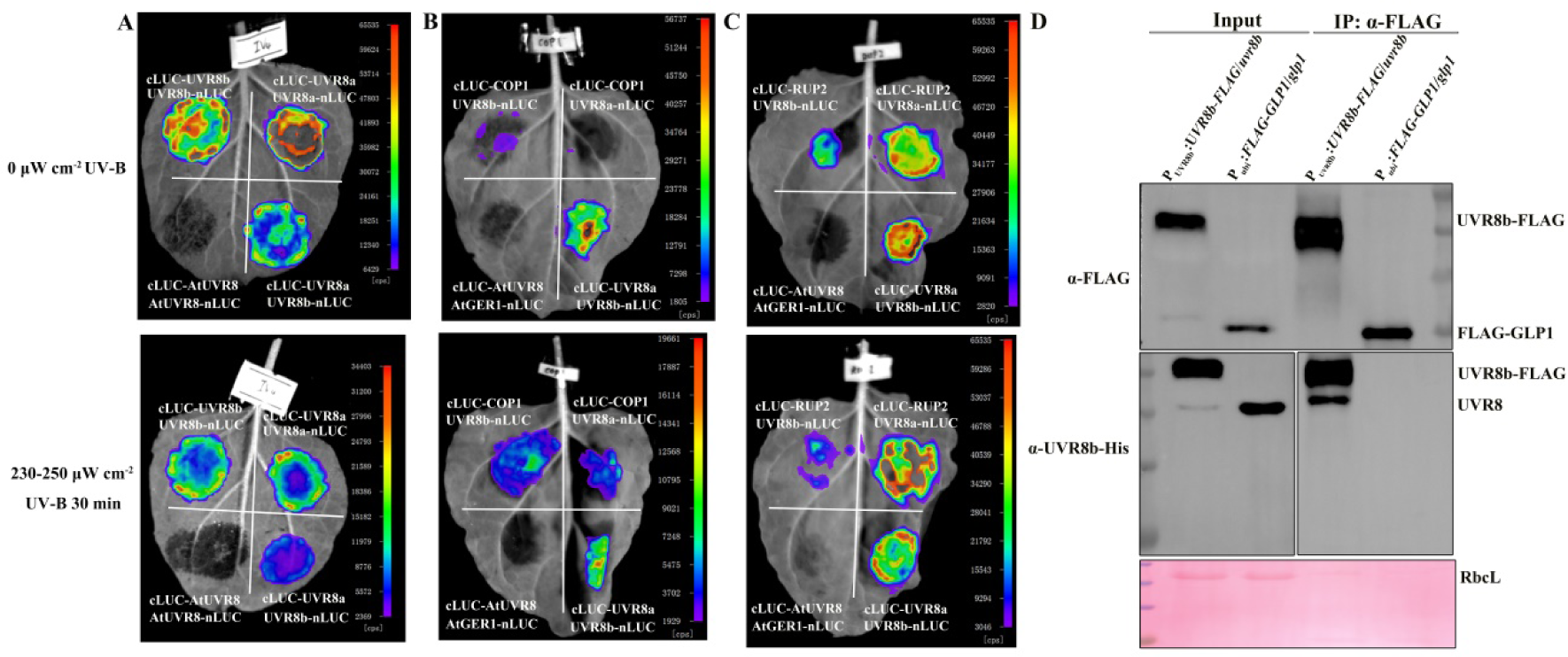
UV-B can affect the interactions of UVR8a/UVR8b and UVR8a, UVR8b, and COP1. Leaf epidermal cells of *Nicotiana benthamiana* were co-transformed with UVR8a-nLUC / UVR8b-nLUC and cLUC-UVR8a (A), cLUC-UVR8b (A), cLUC-COP1 (B) and cLUC-RUP2 (C). D: Co-IP of rice endogenous UVR8a with UVR8b-FLAG from 2-leaf P*_UVR8b_*: *UVR8b-FLAG*/*uvr8b* seedlings transferred from white light (0 μW·cm^−2^ UV-B) to sunlight (55-107 μW·cm^−2^ UV-B) for 2.5 h. Ponceau staining of RbcL is shown as a loading control.

### Overexpression of OsUVR8a/OsUVR8b in rice leads to enhanced UV-B photomorphogenesis and stress tolerance

To further elucidate the role of UVR8 protein in the response of rice to UV-B, transgenic lines overexpressing *OsUVR8a*/*OsUVR8b* under the control of the constitutive strong Ubi promoter were generated. The transgenic lines in which quantitative RT–PCR detected a significantly increased overexpression of *UVR8* mRNA (Figure S7) and Western blot analysis estimated overexpression of UVR8 using anti-rice His-UVR8b were compared with WT in the following detailed analyses (Figure 5A). When grown under sunlight in March for 12 days, the plant height of the *UVR8a*OE and *UVR8b*OE seedlings were lower than those of WT and *uvr8a uvr8b* grown under sunlight (Figure S8A-C). Consistently, mature *UVR8a*/*UVR8b*OE plants grown under sunlight in a paddy field from April to June were also shorter than WT and *uvr8a uvr8b* (Figure S8D and E). However, the seed production rate of the *UVR8a*/*UVR8b*OE plants was slightly lower than that of WT while it was higher than that of *uvr8a uvr8b* (Figure S8F). Plant height of the *UVR8a*OE and *UVR8b*OE seedlings was also lower than those of WT when grown under white light supplemented with narrowband UV-B for 7 days (Figure 5B and C). In addition, UVR8a/UVR8bOE lines showed enhanced UV-B tolerance (Figure 5D) that was even further enhanced upon UV-B acclimation. Therefore, both UVR8a and UVR8b are involved in the UV-B-induced photomorphogenic pathway. Whereas slightly enhanced expression of *HY5* and *RUP2* as well as insignificant *CHS* increase in the leaves of both *UVR8a*OE and *UVR8b*OE seedlings transferred from white light to UV-B were observed (Figure 5 E, F and H), compared to those in WT, suggesting a tight control of the UVR8 in response to UV-B might exist in rice.

**Figure 5.**
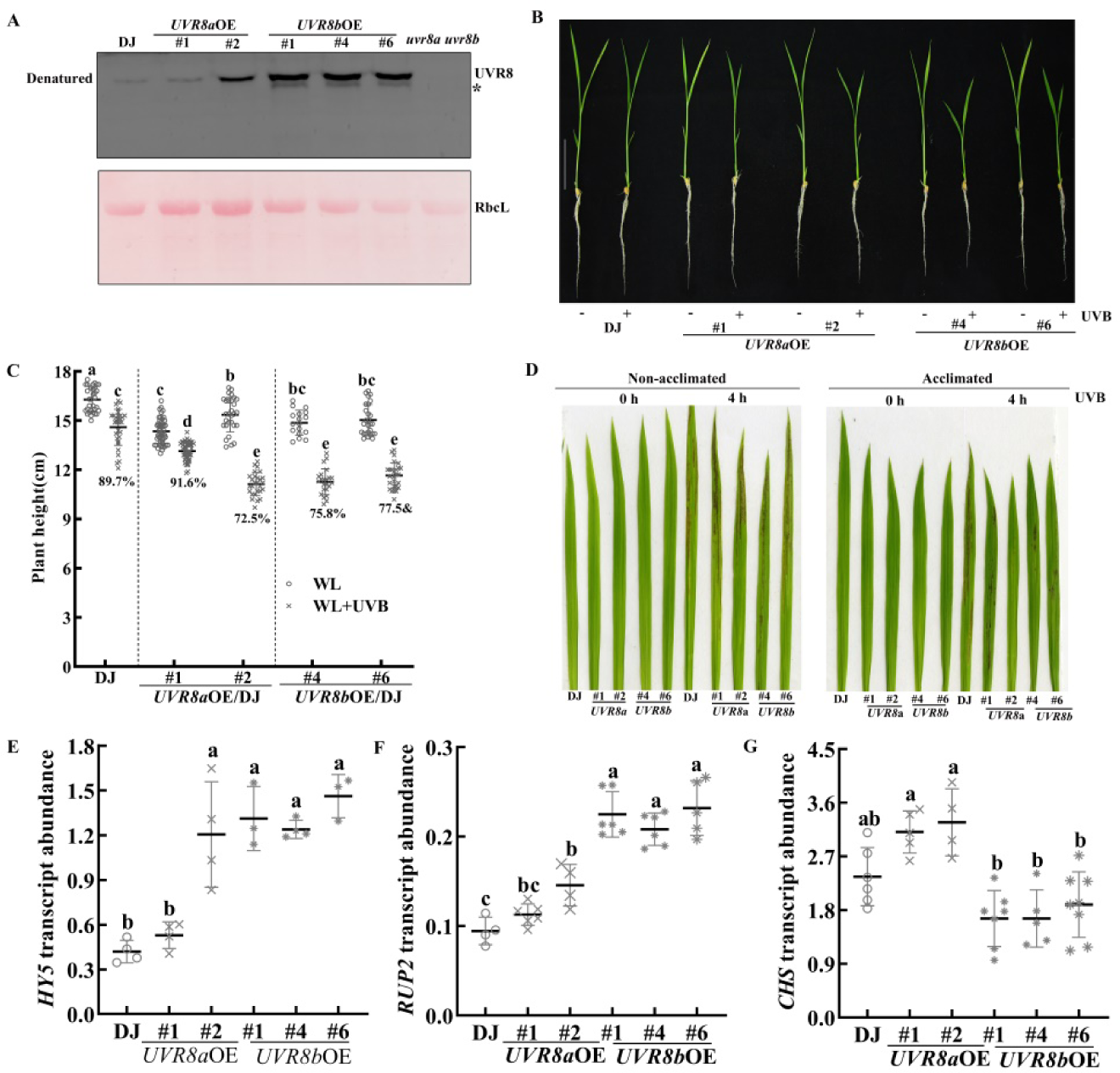
Overexpression of UVR8a/UVR8b enhances UV-B-induced photomorphogenesis and the acclimation of rice to UV-B. A: immunoblot analysis of UVR8 protein levels in the leaves of WT (DJ), *UVR8a*OE (#1, #2), *UVR8b*OE (#1, #4, #6) and *uvr8a uvr8b* seedlings grown under sunlight. Ponceau staining of RbcL is shown as a loading control. B: phenotype of representative seedlings grown for 4 d under dark after imbibition and then exposed to white light or white light supplemented with 54-101 μW·cm^−2^ narrowband UV-B for 7 d. C: quantification of plant height of the seedlings depicted in B. D: phenotype of seedlings grown for 4 d under dark after imbibition and then exposed to white light (non-acclimated) or white light supplemented with 54-100 μW·cm^−2^ narrowband UV-B (acclimated) for 7 d before exposure to 110-130 μW·cm^−2^ UV-B for 5 h. Photos were taken after a 2-d recovery period. E, F and G: quantitative RT-PCR analysis of *HY5* (E), *RUP2* (F) and *CHS* (G) gene expression in response to UV-B in *UVR8a*OE (#1, #2) and *UVR8b*OE (#1, #4, #6) seedlings grown under white light for 8 d after germination and then exposed to 100-110 μW·cm^−2^ UV-B for 4 h compared to WT (DJ). The means assigned with the same letters indicate no statistically significantly difference as determined by one-way different ANOVA with Tukey HSD or Duncan post-hoc multiple comparisons for differences among the means (*P* > 0.05).

## Discussions

### OsUVR8b plays a predominant role in UVR8-mediated UV-B responses

UVR8 is the only UV-B receptor in different plants reported so far (Rizzini et al., 2011), and it has evolved in a conserved manner as a single copy gene in most green plant clades (Zhang et al., 2022). Several species are, however, known to contain at least two UVR8 genes (Brown et al., 2005; Fernández et al., 2016). For example, although the well-known model experimental system, Arabidopsis has one *UVR8* gene, there are two *UVR8* genes in rice. So two rice proteins known as UVR8a and UVR8b exhibit homology to AtUVR8 (Idris et al, 2021). There is, however, a possibility that the different UVR8 gene members, if present, might have distinct roles in UV-B adaptation, at least in rice. In the present study, *UVR8a* and *UVR8b* transcripts could be detected in the leaves, leaf sheaths and roots of WT rice seedlings at the four-leaf stage, but the levels of *UVR8a* and *UVR8b* transcripts and the corresponding proteins were higher in the leaves and leaf sheaths, while the transcript and protein levels of *UVR8b* were higher than those of *UVR8a*. This difference among plant organs could be associated with the necessity of the plant to protect itself from UV-B, and the difference in abundance between UVR8a and UVR8b suggests that OsUVR8b might play a predominant role in rice acclimation to UV-B. In addition, RUP2 exhibited higher interaction intensity with UVR8a than with UVR8b, based on firefly luciferase complementation imaging. This may be due to the difference between UVR8a and UVR8b in their C-terminal sequences as well as structures which play an important role in interaction with COP1 and RUP2 (Figure S1), suggesting that their functions may be different. *uvr8a* mutants exhibited comparable growth inhibition with WT seedlings exposed to specific UV-B radiation and a slight decrease in plant height under paddy field conditions, significant increase in the levels of *CHS* and *HY5* transcripts in the leaves of *uvr8a* mutants transferred to UV-B. Whereas the growth inhibition of *uvr8b* mutant lines was slightly lower than that of WT grown under white light supplemented with narrowband UV-B and there were significant reduced levels of *CHS* and *HY5* gene expression. In addition, plant height, the seed production rate and pollen fertility of *uvr8b* mutants were also significantly lower than those of *uvr8a* mutants and WT grown to the mature stage in a paddy field. *uvr8a uvr8b* mutants, like UV-B light insensitive mutant *uli* and *uvr8*, appeared to be non-responsive to UV-B and a lack of UVB-induced photomorphogenesis as well as *CHS* gene activation. Although the plant height of *uvr8a uvr8b* mutants was insensitive to low-dose UV-B, the mutants were highly sensitive to high-dose UV-B stress, showing phenotypes such as lesion in leaves and reduced acclimation to UV-B, pollen fertility and seed production rate. Moreover, in response to white light supplemented with a high dose of UV-B or in hot summer days, there were more leaf lesions in the leaves of *uvr8a uvr8b* than in *uvr8b* mutants and *uvr8a* mutants, suggesting that UVR8a and UVR8b can participate together in tolerance to UV-B radiation in rice, but UVR8b plays a predominant role in rice acclimation to UV-B. Like Arabidopsis UVR8 overexpressing mutants, they were also more tolerant and adapted better to UV-B, despite no significant UVB-induced an increase in *CHS* transcript levels in OsUVR8a or OsUVR8b overexpressing plants, while plant height of UVR8a or UVR8b overexpressing plants grown under white light supplemented with narrowband UV-B or under solar UVB all displayed enhanced UVB-induced photomorphogenesis. These findings further support that UVR8a and UVR8b can participate together in response to UV-B radiation in rice, but UVR8b plays a prominent role in this process. Nevertheless, significant differences in sequence and tertiary structure are present in the C-termini of OsUVR8a and OsUVR8b, whereas the 27-amino acid region (termed C27: amino acids 397–423) in the C-terminus of UVR8 interacts with COP1, RUPs, WRKY36, BIM1 and BES1 (Cloix et al., 2012, Liang et al., 2018; Yang et al., 2018). However, the precise difference in the participation of two UVR8 genes in rice in response to UV-B remains to be explored further (Figure 6).

**Figure 6.**
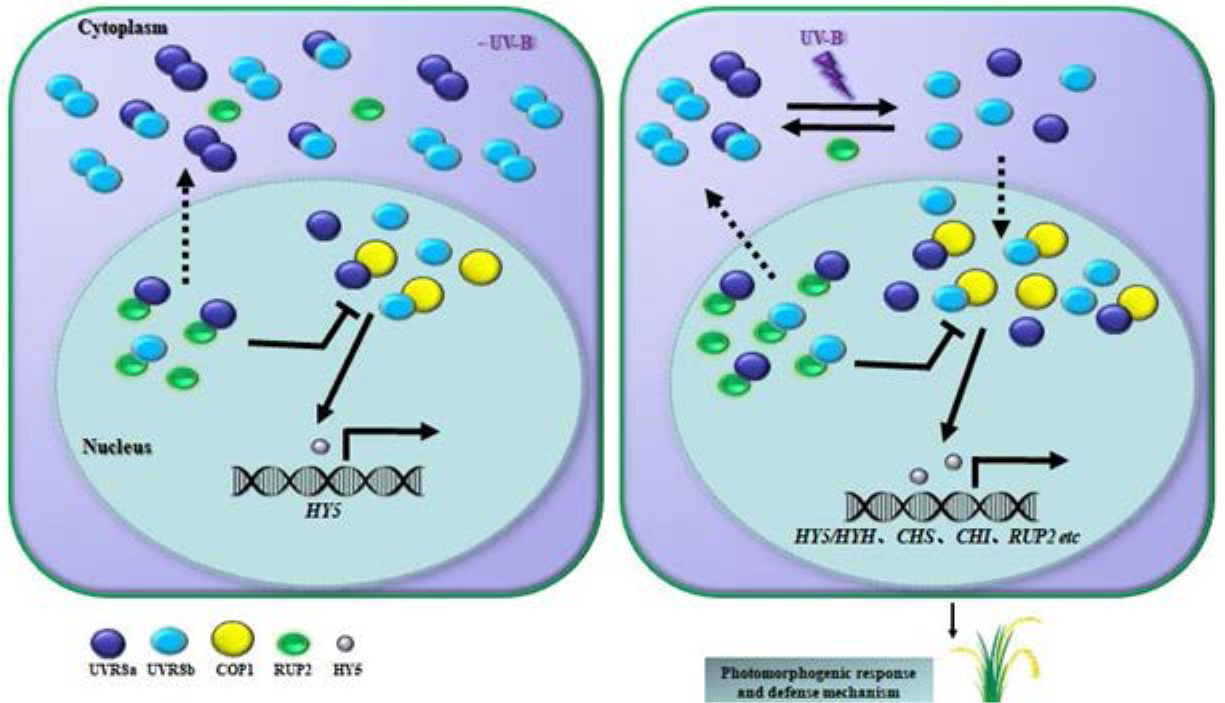
Model of UVR8 actions in rice. In the absence of UV-B, some UVR8 proteins are localized in the cell nuclei and exist as monomers, while the other exists as homodimers or heterodimers in rice. UV-B radiation significantly increases the proportion of the monomer interaction with COP1 rather than the ratio of monomer/dimer. The interactions of the monomers with COP1 and RUP2 initiate transcriptional responses, including stimulation of *CHS*, *HY5* and *RUP2* expression. Purple circles denote UVR8a, blue circles denote UVR8b, yellow circles denote COP1, green circles denote RUP2, gray circles denote HY5, solid lines denote promote and repress, dotted lines denote into and out of the nucleus.

### UVR8a and UVR8b, that have different location and response modes compared to Arabidopsis UVR8, function in rice UVR8-mediated UV-B signaling pathway

UVR8 plays a vital role in promoting UV-B acclimation and tolerance in *Arabidopsis thaliana* by orchestrating the protective gene expression responses that enable its survival under sunlight with high doses of UV-B raditation. The Arabidopsis *uvr8* mutant plants are hypersensitive to UV-B stress (Kliebenstein et al. 2002), as they showed longer hypocotyl, less flavonoid accumulation, and more damage under UV-B radiation than WT and could not survive under sunlight (Brown et al, 2005). On the other hand, overexpression of Arabidopsis UVR8 makes Arabidopsis more tolerant and better adapted to UVB (Martinez et al., 2013). Therefore, UVR8 has been identified as a key positive regulator in UV-B-induced photomorphogenic development and stress acclimation (Brown et al. 2005; Jenkins 2014; Rizzini et al. 2011; Tilbrook et al. 2013). In rice, there are two proteins homologous to AtUVR8, but only one protein with 46% homology to AtRUP2 and no RUP1 sequences have been detected, and rice is adapted to growth in a high light environment with high levels of UV-B. However, it is surprising that the rice *uvr8a uvr8b* double mutants could survive in sunlight with slightly decreased plant height, pollen vitality and seed production rate, given that the conserved amino acids in OsUVR8a / OsUVR8b are same as those in AtUVR8, moreover, UV-B-induced activation of *CHS* and *HY5* depends on OsUVR8a and OsUVR8b, and OsUVR8a/OsUVR8b can interact with COP1 and RUP2. These findings suggest that other mechanisms independent of UVR8-induced UV-B acclimation might operate in rice. In Arabidopsis grown under white light, UVR8 predominantly exists as a dimer and is localized in the cytosol. Upon UV-B photoreception, stimulation of nuclear accumulation of the UVR8 was observed in Arabidopsis, and the dimer then rapidly dissociated into monomers which interacted with COP1 into the nucleus (Heijde and Ulm, 2013; Kaiserli and Jenkins, 2007; Rizzini et al, 2011). GFP fluorescence of OsUVR8a/OsUVR8b was present in the nucleus even if the protoplasts were isolated from etiolated seedlings or tobacco grown in minus UV-B (Figure S6) and part of OsUVR8a/OsUVR8b proteins exist as monomers in the absence of UV-B, the ratio of OsUVR8 protein dimer/monomer showed insignificant change in the leaves of rice seedlings transferred from white light to white light supplemented with UV-B. These might make rice help to grow in high level UV-B, because UV-B-acclimated *Arabidopsis* exposed to 15-fold increase in UV-B also did not exhibit significant increase in the level of monomer although UVR8 can mediate the response to UV-B (Liao et al, 2020). Additionally, fluorescence signals of OsUVR8a-nLUC and cLUC-OsUVR8b along with OsUVR8b-nLUC and cLUC-OsUVR8a/cLUC-OsUVR8b still keep high level in firefly luciferase complementation experiments even if the tobacco exposed to 230-250 μW·cm^−2^ UV-B, moreover, OsUVR8a can interact with OsUVR8b under sunlight with about 100 μW·cm^−2^ UV-B, indicating OsUVR8a and OsUVR8b can exist as heterodimer except homodimer even if rice exposed to UV-B (Figure 4 and 6). These differences might attribute to small variations in amino acid sequences outside the conserved motifs. Such as the UVR8 from *M. polymorpha* is also mainly present as a monomer in absence of UV-B and constitutively in the nucleus even if functional motifs that regulate protein stability and localization are conserved (Soriano et al, 2018). Besides, differences between AtUVR8 and OsUVR8 sequences as well as tertiary structures mainly exist in N-terminus and C-terminus (Figure S1), whereas 23-aa region in the N-terminus of AtUVR8 is required for efficient nuclear accumulation (Kaiserli and Jenkins, 2007) and 27-aa region in the C-terminus interacts with COP1, RUPs, WRKY36, BIM1 and BES1 (Cloix et al., 2012; Liang et al., 2018; Yang et al., 2018).The interaction between COP1 and UVR8 enhances the accumulation of HY5 and activates the expression of a set of genes (Yin et al, 2016), therefore, HY5 is a key effector of the UVR8 pathway, and that it is required for the survival of Arabidopsis under UV-B radiation (Brown et al. 2005). In Arabidopsis, UV-B–induced expression of the *HY5* gene was impaired in *uvr8* mutant and enhanced in UVR8 overexpressing plants (Brown et al, 2005), but the transcript level of *HY5* in rice that is homologous to *AtHY5* showed a slight increase and insignificant *CHS* change in the leaves of OsUVR8a or OsUVR8b overexpressing plants exposed to UV-B compared with those of in WT. These suggest that a more rigorous UVR8-mediated UV-B signaling regulatory mechanism might exist in rice. Moreover, only about 30% of UVB-regulated genes of rice were found to have Arabidopsis homologs, while a large portion of the regulated rice genes had no homologs reported to be UV-B-regulated in Arabidopsis (Idris et al, 2021). In addition, unlike *AtUVR8*, the transcript levels of *UVR8a* and *UVR8b* were downregulated in the leaves of rice seedlings transferred from minus to increase in UV-B, this is same with the reports about homologies of *AtUVR8* in tea and *Brachypodium distachyon* response to UV-B (Shamala et al., 2020; Chen et al, 2023). Therefore, although UVR8-mediated signaling pathway is conservative in rice, but OsUVR8 location and the mechanism of response to UV-B as well as the downstream regulators and transcription factors might be different from those of Arabidopsis. UV-B activates UVR8-mediated UV-B responses in rice might depend on increasing the proportion of monomers bound to COP1 to cause changes in downstream genes, rather than promoting an increase in monomer abundance and entry into the nucleus (Figure 6). Moreover, it remains to be determined if other types of UV-B receptors might be found in rice.

## Materials and Methods

### Plant materials and growth conditions

Rice (*Oryza sativa* L.) cv. Dongjin (DJ) was used for generating the *uvr8a*, *uvr8b*, *uvr8a uvr8b* double mutants and transgenic rice plants overexpressing *OsUVR8a* / *OsUVR8b*. Germinated seeds were grown in the Kimura B complete nutrient solution (Yoshida et al., 1976) under different experimental conditions as described in the legend of each figure. A plant growth chamber (PERCIVAL E-41HO) was fitted with fluorescent lamps to supply white light. For supplementary UV-B in some experiments, Philips TL20W/01 UV-B tubes (Germany, with a radiation spectrum of 300-320 nm and a peak at 311 nm) were used for narrowband UV-B irradiation, while Sankyo G15T8E UV-B tubes (Japan, with a radiation spectrum of 290-310 nm and a peak at 306 nm) were used in other experiments. The UV light intensity of the lamp was measured using a 742 UV light meter (manufactured by the Beijing Normal University, Beijing, China).

### Generation of transgenic rice plants

The *uvr8a*, *uvr8b*, *uvr8a uvr8b* mutants were generated using the CRISPR/Cas9 system. Firstly, the vector containing a CRISPR cassette comprising a functional Cas9 under an Ubi promoter and gRNA with targets at different locations of *OsUVR8a* or/and *OsUVR8b* was constructed according to Ma et al. (2015). The vector was then introduced into the *Agrobacterium tumefaciens* strain EHA105 for transformation of DJ rice calli using the procedure as described in Hiei et al. (1994). The transgenic plants harboring the mutations in *OsUVR8a* or/and *OsUVR8b* were regenerated from the transformed rice calli and were then was genotyped by PCR amplification of the target region in DNA extracted from their leaves. The primer sequences used for vector construction and amplification of the target region are listed in the Supplementary Table S1 and Table S2. For *OsUVR8a*OE/*OsUVR8b*OE vector construction, the full length *OsUVR8a*/*OsUVR8b* coding sequence was amplified using PCR with the primers listed in the Supplementary Table S2 and inserted into the pOx vector (provided by Professor Yao-Guang Liu, College of Life Sciences, South China Agricultural University, China) containing a maize ubiquitin (Ubi) promoter. The *OsUVR8a*OE/*OsUVR8b*OE vector was introduced into the *Agrobacterium tumefaciens* strain EHA105 for transformation of DJ rice calli according to Hiei et al. (1994).

### Total RNA extraction and RT-qPCR analysis

Total RNAs were extracted from various tissues of wild-type (WT) plants or *OsUVR8* transgenic plants with Trizol and reverse-transcribed with HiScript® II following the manufacturer’s instructions (Vazyme Biotech, Nanjing, China). RT-qPCR analyses were carried out using a PTC200 (BIO-RAD) PCR machine and a SYBR green probe (Bimake, China). The primers used are given in the Supplementary Table S2. ACTIN was used as an internal standard.

### Subcellular localization of OsUVR8a and OsUVR8b

The full-length coding region of OsUVR8a or OsUVR8b was amplified by PCR using rice leaf cDNA as a template. The PCR products were cloned into the vector pYL322-d1-GFP or pCAMBIA1300-(N)eGFP to produce a fusion protein with green fluorescence protein (GFP) under the control of the 35S promoter. Rice protoplast isolation and PEG-mediated transient expression of pYL322-d1-GFP/UVR8a-GFP/UVR8b-GFP were carried out according to Zhang et al (2011). Transformation of *Nicotiana benthamiana* expressing pCAMBIA1300-GFP/GFP-UVR8a/ GFP-UVR8b was carried out according to Qi et al (2016). Green / red fluorescence from the expression of GFP / RFP in rice protoplast was observed using an argon laser and the PMT detectors of Zeiss LSM 7 DUO. The excitation/emission filters utilized for fluorescence detection were 488/493-546 nm for GFP, and 532/570-620 nm for RFP. Green from the expression of GFP in *Nicotiana benthamiana* was observed using Leica DFC550.

### Firefly luciferase complementation fragment imaging assay

The gene constructs (cLUC and nLUC) for fusion proteins between the C or N terminus of firefly luciferase and UVR8a, UVR8b, COP1, or RUP2 was transferred to the *Agrobacterium* strain GV3101. The *Agrobacterium* strains carrying the different constructs were simply mixed and infiltrated into leaves of *N. benthamiana*. The transformed tobacco plants were kept under long-day white light for 36-48 h. A luciferin solution (1 mM luciferin and 0.01% Triton X-100) was then infiltrated into the leaves co-expressing the different constructs and adapted in the NightSHADE LB98 Imaging System for 5 min before the leaves were examined for LUC activity (Chen et al, 2008). Images were captured using the NightSHADE LB98 imaging system (Berthold Technologies, Bad Wildbad, Germany).

### Protein gel blot and Co-immunoprecipitation analysis

Leaves, leaf sheaths and roots of rice seedlings were ground in liquid nitrogen, and the proteins were extracted with a buffer consisted of the following: 50 mM Tris-HCl pH 7.5, 150 mM NaCl, 1 mM EDTA, 0.5% (v/v) Triton X-100, 1 mM PMSF (BBI, product no. A610425), 10 μM proteasome inhibitor MG132 (Sangon Biotech, product no. T510313, Shangai, China) and 1% (v/v) protease inhibitor cocktail for plant extracts (Sigma, product no. P9599). The supernatant was mixed with loading buffer, and was then separated using 10% SDS-PAGE electrophoresis. After electrophoresis, the gels were irradiated with UV-B light (95-100 μW·cm^−2^ for 15 min), then the proteins were transferred to Nitrocellulose (NC) / PVDF membranes. Anti-His-OsUVR8b prepared in our laboratory at 1: 1000 dilution or anti-FLAG VHH HRP (AlpalifeBio^TM^, product no. KTSM1318, Shenzhen, China) at 1:5000 dilution was used as the primary antibodies. Horseradish peroxidase (HRP)-conjugated anti-rabbit (BBI D110011) was used as the secondary antibody. Immunodetection was performed using an ECL Enhanced Plus Kit (ABclonal, Wuhan. China) and detected using a ChemiScope 6000 Touch chemiluminescence imaging system (Clinx). For immunoprecipitation analysis, 2-leaf stage P*_UVR8b_*: *UVR8b-FLAG*/*uvr8b* and P*_Ubi_*: *FLAG*-*OsGLP1*/*glp1* rice seedlings grown under white light (0 μW·cm^−2^) were transferred to sunlight (55-107 μW·cm^−2^) for 2.5 h, then whole seedlings were ground in liquid nitrogen, and the proteins were extracted with the above-mentioned buffer. The supernatant was mixed with 10 μL ANTI-FLAG^®^M2 Magnetic beads (Sigma, product no. M8823), incubated at 4℃ for 5 h, and then the beads were washed three times with TBS (50 mM Tris-HCl pH 7.5 containing 150 mM NaCl). The bound proteins were eluted from the affinity beads with 2×SDS loading buffer at 95℃ for 10 min and the supernatant were used for protein gel blot analysis.

## Supplementary Data

**Supplementary Figure S1** Differences in amino acid sequence and tertiary structure among OsUVR8a, OsUVR8b and AtUVR8 were found mainly in the N- and C-termini.

**Supplemental Figure S2** Immunoblot analysis of UVR8 protein with anti-His-OsUVR8b in the leaves of Arabidopsis Col-0 seedlings exposed to UV-B radiation for 0 h, 1 h, 2 h and 4 h.

**Supplemental Figure S3** UV-B stimulates the expression of *HY5*, *RUP2* and *CHS*.

**Supplemental Figure S4** Sequences of sgRNA target region of WT, *uvr8a*, *uvr8b* and *uvr8auvr8b*.

**Supplemental Figure S5** Mutation of *UVR8a* or / and *UVR8b* impairs the tolerance to UV-B of rice.

**Supplemental Figure S6** OsUVR8a/OsUVR8b GFP-fusion proteins were localized in the nucleus even in the absence of UV-B.

**Supplemental Figure S7** Molecular evaluation of transgenic plants overexpressing *OsUVR8a*/*OsUVR8b*.

**Supplemental Figure S8** Overexpression of UVR8a/UVR8b exacerbates UV-B-induced growth inhibition of rice.

**Table S1** Primer sequences used for vector construction

**Table S2** Primer sequences used for qRT-PCR and amplification of the target region

## Acknowledgments

This work was supported by the National Natural Science Foundation of China (Grant No. 32171934) and Natural Science Foundation of Guangdong (Grant No. 2020A1515010192). We thank Yaoguang Liu and Qinlong Zhu (South China Agricultural University) for kindly providing pOX and CRISPR/Cas9 system vectors, Huili Liu and Hai Zhou (South China Agricultural University) for kind help and helpful advice, Lili Cui and Xiangyang Li for kind help in firefly luciferase complementation fragment imaging assay, Xiaojing Zhang (South China Agricultural University) for GFP fluorescence observation.

## Author contributions

Chen Y, Zhong Y, Yan X, Peng X and Liu E designed the research. Chen Y, Zhong Y, Yan X, Ouyang M, Ye Y, Li S and Liu E performed the experiments, analyzed the data. Leung DWM, Chen Y, Zhong Y, Peng X and Liu E wrote the manuscript. Chen Y and Zhong Y contributed equally.

